# Morphological and genomic characterisation of the hybrid schistosome infecting humans in Europe reveals a complex admixture between *Schistosoma haematobium and Schistosoma bovis parasites*

**DOI:** 10.1101/387969

**Authors:** Julien Kincaid-Smith, Alan Tracey, Ronaldo Augusto, Ingo Bulla, Nancy Holroyd, Anne Rognon, Olivier Rey, Cristian Chaparro, Ana Oleaga, Santiago Mas-Coma, Jean-François Allienne, Christoph Grunau, Matthew Berriman, Jérôme Boissier, Eve Toulza

## Abstract

Schistosomes cause schistosomiasis, the world’s second most important parasitic disease after malaria. A peculiar feature of schistosomes is their ability to produce viable and fertile hybrids. Originally only present in the tropics, schistosomiasis is now also endemic in Europe. Based on two genetic markers the European species had been identified as a hybrid between the ruminant-infective *Schistosoma bovis* and the human-infective *Schistosoma haematobium.*

Here we describe for the first time the genomic composition of the European schistosome hybrid (77% of *S. haematobium* and 23% of S. *bovis* origins), its morphometric parameters and its compatibility with the European vector snail and intermediate host Compatibility is a key parameter for the parasites life cycle progression. We also show that egg morphology (a classical diagnostic parameter) does not allow for differential diagnosis while genetic tests do so. Additionally, we performed genome assembly improvement and annotation of *S. bovis,* the parental species for which no satisfactory genome assembly was available.

For the first time since the discovery of hybrid schistosomes, these results reveal at the whole genomic level a complex admixture of parental genomes highlighting (i) the high permeability of schistosomes to other species’ alleles, and (ii) the importance of hybrid formation for pushing species boundaries not only conceptionally but also geographically.

## Author summary

In 2013, schistosomiasis reached Southern Europe. Since then, endemic infections were recurrently identified in 2015 and 2016, clearly indicating that the parasite has settled and established locally. Using two molecular markers, we had previously demonstrated that the parasite is a hybrid between *Schistosoma haematobium* and *S. bovis* that are known to infect humans and livestock, respectively. Nevertheless, this method had very low resolution and did not allow to give clear answers on the origins and the mechanisms of hybrid generation, e.g. if the hybrid had been generated recently on Corsica or if it invasive. The genome-wide sequencing approach used in this work allowed us to reveal a complex admixture between the parental genomes and suggests that hybridization between these two species may be the result of ancient crossing events.

Additionally, wherever in Africa or in Europe, a clear discrepancy exists between the egg shape usually used for species identification and the genomic composition. Therefore, egg shape cannot be used anymore as a good indicator for hybrid detection. Knowing the traits and the genetic features of the hybrid has implications in terms of diagnostic as well as disease management either through vector control strategies or treatment of patients.

## Introduction

Schistosomes are parasitic flatworms, responsible for the major tropical disease schistosomiasis, The epidemiological statistics associated with the disease are sobering: 800 million people are at risk in 78 countries, mostly concentrated in sub-Saharan Africa; 230 million are infected and the disease causes more than 200 000 deaths each year as well as between 1.7 and 4.5 million Disability Adjusted Life Years (DALYs) [1]. The most exposed groups are children and young adults that have predominant activities linked to contaminated freshwater environments. In addition to humans, schistosomiasis severely impacts livestock in Africa and Asia where an estimated 165 million animals are infected [2].

Schistosomes have a complex life cycle that includes passage through a freshwater snail intermediate host (hereafter designed as the parasite vector) and a final vertebrate definitive host. Global changes, both anthropogenic and environmental modifications, may contribute to modifications in the geographical distribution of species and expand their potential ecological niches [3]. Distinct species may thus acquire a new capacity to interact, hybridize and subsequently introgress their genomes by backcrossing with parental species or other hybrids, a phenomenon called “hybrid swarm”. Hybridization between individuals from two previously reproductively isolated species is generally expected to produce offspring less fit than the parents, sometimes infertile or even sterile but in some cases however, hybridization may lead to greater fitness than parental species, also known as hybrid vigour or heterosis [4], Hybridizations of human parasites are not scarce and are a real concern in terms of parasite transmission, epidemiology and disease [5].

Hybridizations between schistosomes have already been identified: (i) between different human-specific schistosome species, (ii) between different animal-specific schistosome species, (iii) and between human-specific and animal-specific schistosomes species [6]. These latter hybrid forms are particularly alarming because they raise the possibility of the emergence of new zoonotic parasitic strains, introducing an animal reservoir, and therefore greatly hampering our ability to properly control the life cycle. The precise characterization of the introgression levels of hybrid populations is essential for diagnostics purpose but is also necessary to better understand the parasite life history traits, the disease dynamics and the epidemiology in the field. Next generation whole-genome sequencing is now the tool of choice for a deeper insight into the genomic composition of natural hybrids.

In this study, we aimed at fully characterize schistosome hybrids that have recently emerged in Europe. We described hybrids based on the morphology of eggs, the disease- and diagnostic-relevant stage, as well as compatibility with potential European snail vectors. We characterized the extent of introgression in natural hybrids at the whole genome scale.

## Results

### Morphology of the eggs does not allow for differential diagnosis of the hybrid

Morphometric analysis is a classical way to identify species. In Appendix S1 we deliver a precise morphological description of the adults and eggs of the European hybrid. Since eggs are easily accessible in the field, they are commonly used to diagnose the infecting schistosoma species. Therefore, egg morphology was of particular interest to us. A total of 44 eggs collected from hamsters infected with the European hybrid were examined for morphological characterization. The length and width were assessed on all eggs, but the spine length has been measured on a subset of 36 that had a spine distinctive enough to allow for proper estimation of their sizes. The results are presented in Table 1.

**Table 1:**
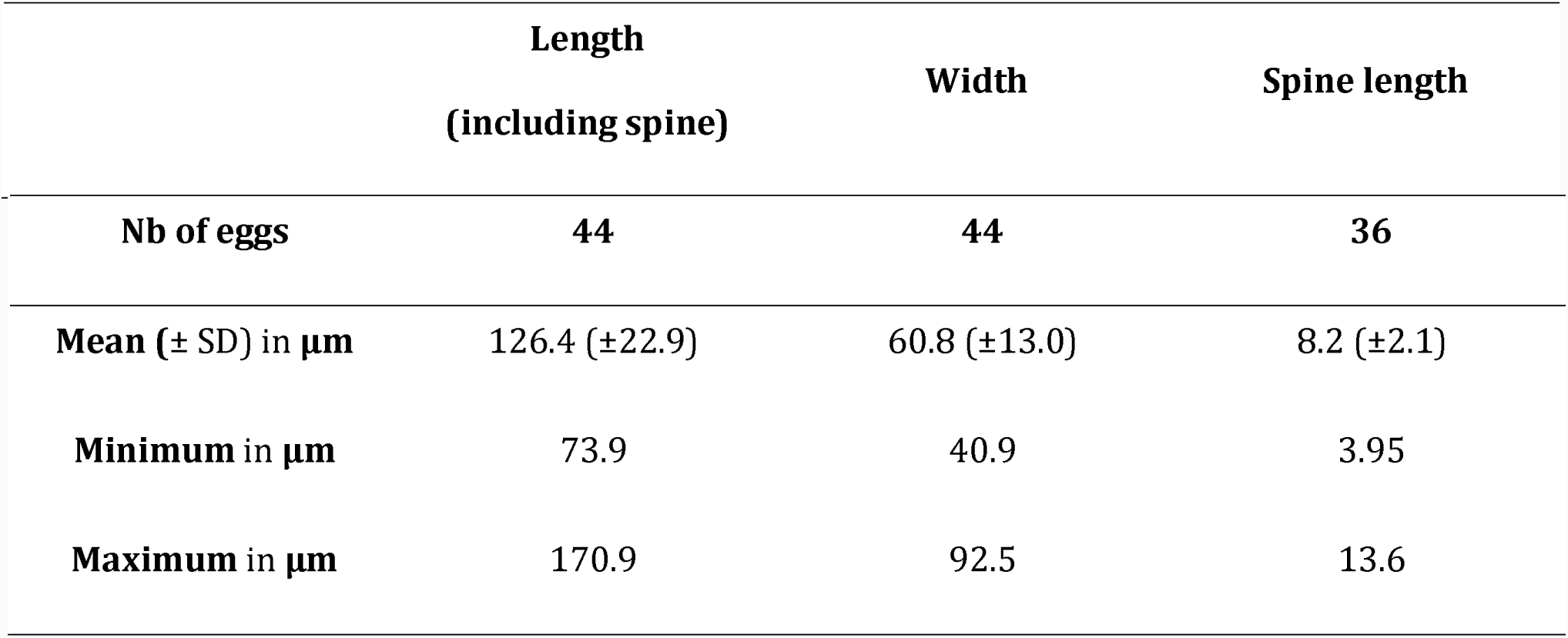
Morphological measurements results of the Corsican hybrid schistosome eggs (SD = Standard deviation).

Most of the eggs displayed a representative elliptical morphotypes and were characterized by a terminal spine, reminiscent of *S. haematobium* infection in humans. Nevertheless, not all eggs had a typical *S. haematobium* morphotype and were in some cases intermediate with *S. bovis-*type eggs (Fig 1). In summary, egg morphology alone cannot differentiate between the hybrid and putative parental species. Nevertheless, this morphological analysis supported the initial conclusion, based on only two genetic markers, that the hybrid originates from a cross between *S. bovis* and *S. haematobium.* **Whole genome sequencing shows introgression of *S. bovis* into *S. haematobium*** To precisely characterize the level of hybridization we anticipated to use Illumina short read massive sequencing and alignment to parental genomes. As the previous *S. bovis* assembly was highly fragmented (111 328 scaffolds, N50 7kb), we resequenced the genome using PacBio long reads. This produced 4,102,584 filtered subreads (48,987,175,429 bases). Genome assembly led to a new genome version of 486 scaffolds (N50 3.1Mb). Gene prediction led to identification of 14,104 protein-coding genes of 18,725 bp average length (5.3% of genome length) (Table 2) which is consistent with the known characteristics of *Schistosoma* genomes [7],

**Table 2:** Assembly and gene prediction metrics of *S. bovis.*

|  |  |
| --- | --- |
| Proportion of genome that is coding (%) | 5.27% |
| Number of genes | 14,104 |
| Average gene length (bp) | 18,725 ± 21,625 |
| Average number of exons per gene | 7.9 |
| Average exon length (bp) | 299 |
| Average intron length (bp) | 2,389 |
| Repeat rate (%) | 49.5% |

**Figure 1:**
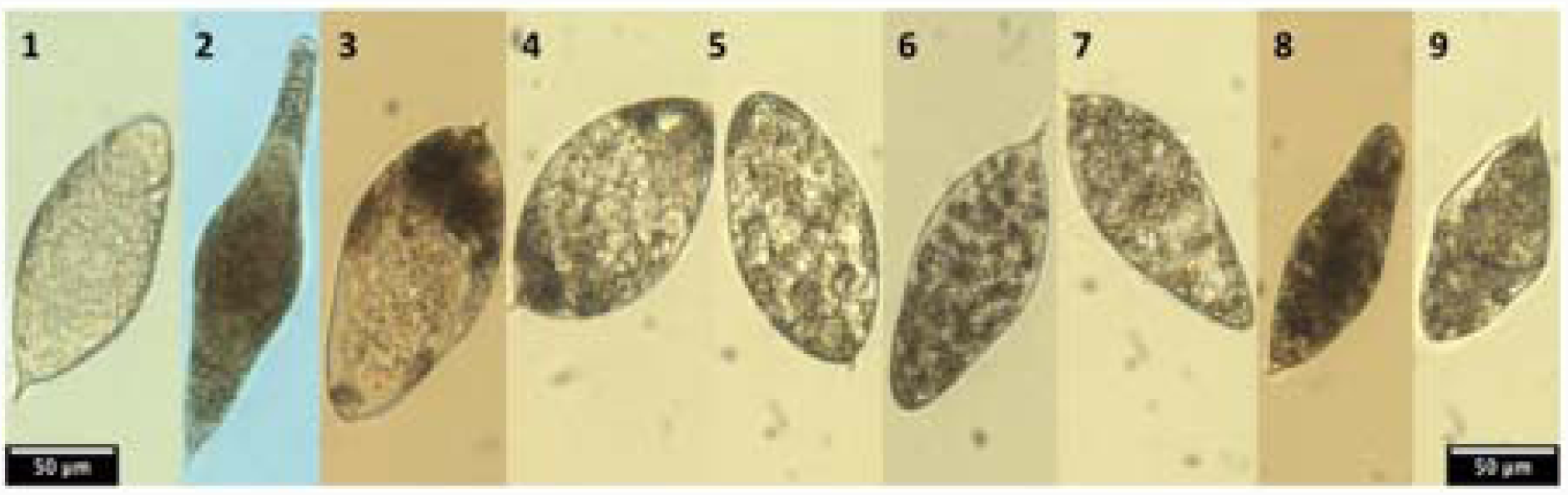
Egg morphologies of the pure parental species and the Corsican *S. haematobium X S. bovis* hybrid. Eggs 1 and 2 show typical morphologies of *S. haematobium* (elliptical with a terminal spine) and *S. bovis* (spindle shape with a terminal spine), respectively. Eggs 3-8 show the egg morphology of the Corsican hybrid schistosome. While most eggs were typical to *S. haematobium* (3-7), a high variability of morphotypes was observed (8-9).

For the European hybrid, a total of 289,873,531 reads (76.5%) were mapped against the 681.2 Mb concatenated genomes of *S. haematobium* [7] and *S. bovis* (this work), representing 42X mean coverage (Table 3). We also sequenced F1 males from experimental first generation cross between male *S. haematobium x* female *S. bovis* as a control. A total of 4,910,354 reads (92.3%) were mapped against the concatenate of *S. haematobium* and *S. bovis* genomes.

**Table 3:**
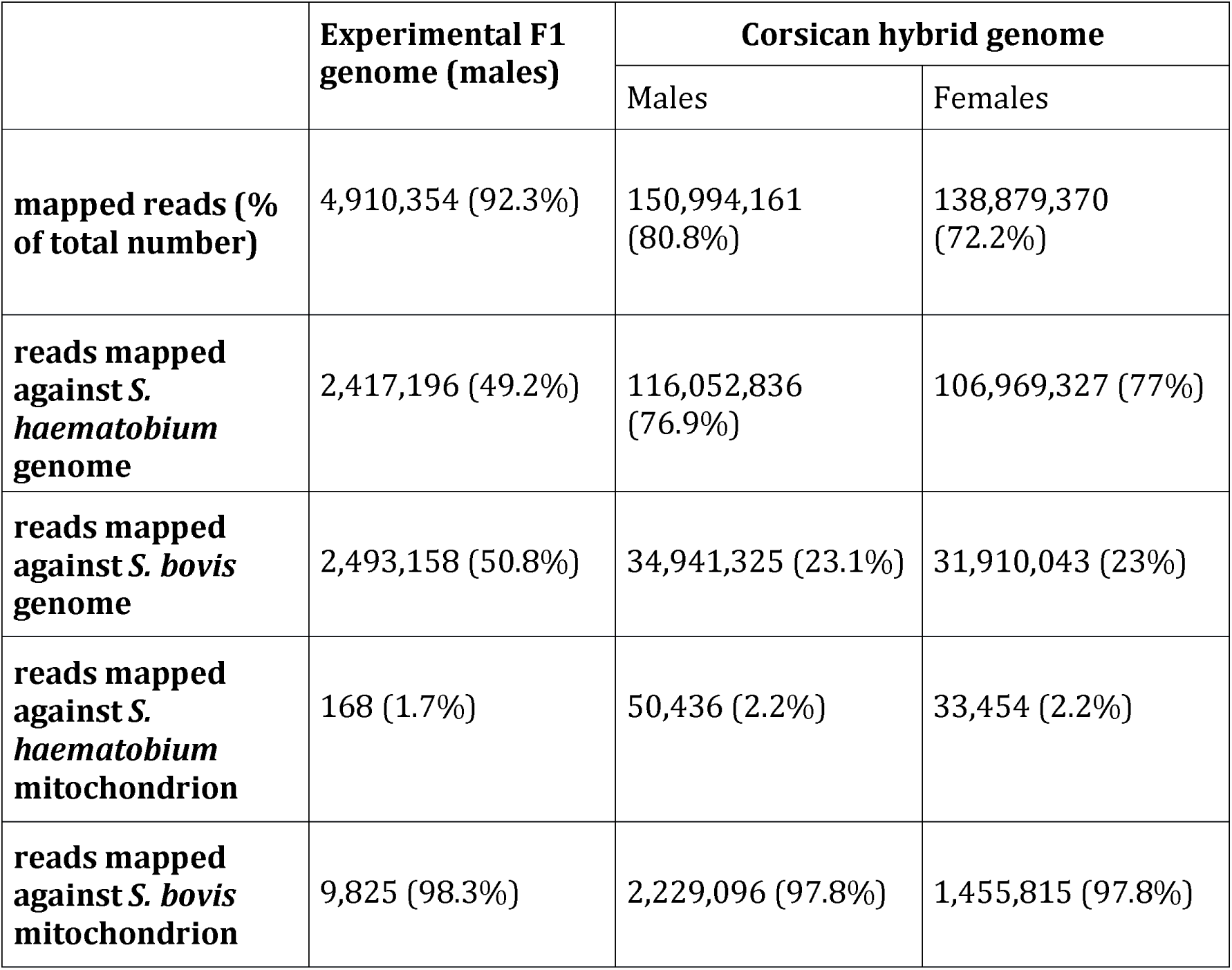
Summary of the introgression level analysis of the experimental F1 hybrids (control) and the Corsican natural hybrid strain.

|  | Experimental F1<br>genome (males) | Corsican hybrid genome |  |
| --- | --- | --- | --- |
|  |  | Males | Females |
| **mapped reads (%**<br>**of total number)** | 4,910,354 (92.3%) | 150,994,161<br>(80.8%) | 138,879,370<br>(72.2%) |
| **reads mapped**<br>**against *S.***<br>***haematobium***<br>**genome** | 2,417,196 (49.2%) | 116,052,836<br>(76.9%) | 106,969,327 (77%) |
| **reads mapped**<br>**against *S. bovis***<br>**genome** | 2,493,158 (50.8%) | 34,941,325 (23.1%) | 31,910,043 (23%) |
| **reads mapped**<br>**against *S.***<br>***haematobium***<br>**mitochondrion** | 168 (1.7%) | 50,436 (2.2%) | 33,454 (2.2%) |
| **reads mapped**<br>**against *S. bovis***<br>**mitochondrion** | 9,825 (98.3%) | 2,229,096 (97.8%) | 1,455,815 (97.8%) |

As expected, the mapping of the FI reads to *S. haematobium* and *S. bovis* genomes gave a proportion that matched the expectations for a first generation hybrid (~50% of each of the parental species genomes; Table 3). This control was crucial to validate our analytical pipeline and thus the results obtained for the Corsican natural European hybrid parasite. Moreover, 98.3% of reads of mitochondrial origin aligned to *S. bovis,* which is consistent with maternal inheritance of the mitochondrial genome.

Interestingly, the mapping of the Corsican hybrid reads against *S. haematobium* and *S. bovis* reference genomes revealed a complex admixture between the parental genomes with a proportion of 76.9% of sequences mapping on *S. haematobium,* and 23.1% mapping on *S. bovis* genomes for both male and female parasites (Table 3). Alignment to the mitochondrial genomes of both parents also showed results concordant with the previous Sanger sequencing data for this marker, with 97.8% of reads being mapped on *S. bovis* mitochondrial genome, and 2.2% on *S. haematobium* mitochondrial genome [8]. To figure out the divergence level between the two pure species genomes, we identified the orthologous regions between *S. haematobium* and *S. bovis* using CACTUS. A total of 234,5 Mb sequences aligned between the parental species, which represents 64% of the *S. haematobium* genome length. The mean similarity of these shared sequences was 95.88%.

Besides being of fundamental interest for the evolutionary biology of the parasite, this finding could also have immediate consequences for parasite control. One of the few phenotypic features that are of relevance for infection success and for which the genetic basis is known is resistance to Oxamniquine (OXA). In *S. mansoni* the mutations that confer resistance occur in the SmSULT-OR gene (Smp_089320), encoding a sulfotransferase that is required for drug activation, are p.E142del and p.C35R [9,10]. In *S. haematobium* the drug is predicted not to be efficient due to a F39 *Sm* > Y54 *Sh* substitution [9]. We used blast searches with Smp_089320 against the *S. haematobium* and the new *S. bovis* genome to identify orthologues. Orthologues exist in both genomes and have no mutations in *Sm* C35 or *Sm* L256 or deletion in *Sm* E142, but both possess the F39 > Y54 which is predicted to negatively impact drug binding. We expect therefore the European hybrid to be genetically resistant to OXA, excluding the drug as treatment option.

We conclude that the European hybrid was generated by an initial cross between *S. haematobium* and *S. bovis* and successive backcrosses with *S. haematobium.*

### The European hybrid is compatible with the natural hosts of *S. haematobium, Bulinus truncatus* but not with *Planorbarius metidjensis,* host of *S. bovis*

The presence of 23% of *S. bovis* genetic information in the hybrid genome raised the possibility that the hybrid was capable of infecting the vector snail of *S. bovis.* To test this hypothesis we exposed *Planorbarius metidjensis,* which is a specific host to *S. bovis,* to miracidia of the natural Corsica hybrid The susceptibility of the parental snail hosts was inferred at 35 days after exposition to 5 parasite larvae (miracidia). The prevalence in *Bulinus truncatus* was 24% (9 infected snails out of 37 alive). In *Planorbarius metidjensis* the prevalence was 0% (0 infected snails out of 29 alive), but we cannot exclude the possibility that other strains of *P. metidjensis* could be compatible with the European schistosome hybrid.

### Discussion

The emergence of new infectious diseases and pathogens is currently among the great concerns of our changing world and has strong outreach effects for society. Besides the important impacts that global changes (climate changes and anthropogenic activities) may have on the spread and transmission of tropical infectious diseases in higher latitudes, other phenomenon may combine and act as a driving force promoting the emergence of novel disease in unsuspected areas. The importance and the frequency of hybridization in infectious agents are certainly underestimated, and very little attention has been given so far to the role of gene introgression on infectious disease emergence, spread and control [11]. In the genus *Schistosoma,* latest reports have revealed that hybrids are frequent throughout West Africa, and are already a real concern for human health [5,6]. Nevertheless, it is the first time that a hybrid schistosome is involved in a large-scale outbreak in Europe [8,12,13]. Although usually restricted to tropical areas, schistosomiasis is now persisting in Corsica and the hybrid status of the parasite might have increased its invasive and adaptive capacities. The hybrid status of the parasite may indeed have important implication for disease control in term of host spectrum, diagnostic, and treatment.

### Implications for host spectrum and parasite distribution

*S. haematobium* and *S. bovis* have different intermediate host specificities. *S haematobium* only infect mollusks from *Bulinus* genus while *S. bovis* also infect *Planorbarius* snails (widely present in the Iberian Peninsula). The potential distribution range of the disease may be enhanced if the hybrid is able to infect the intermediate hosts of both parental species. Interestingly, the natural Corsican hybrid schistosome that we recovered was not able to infect our laboratory strain of *Planorbarius metidjensis,* but displayed high levels of compatibility with *Bulinus truncatus* from Corsica (24% of prevalence) which is consistent with previous compatibility assessments for the hybrid parasite recovered directly from an infected patient in 2014 [8]. This also goes in line with the predominance of *S. haematobium* sequences in this hybrid, and may open the door for vector based control, e.g. by outcompeting these snails. One other fundamental concern is the capacity of such introgressed schistosomes to infect livestock or other reservoir hosts. The zoonotic potential of the hybrids would strongly impact the parasite transmission in the field in and out of endemic areas, and may hamper our capacity to maintain adequate control strategies as schistosomiasis treatment focuses almost exclusively on humans. Nevertheless, despite a recent study showing the presence of pure species (*S. bovis, S. haematobium,* and *S. mansoni)* as well as hybrids between *S. bovis* and *S. haematobium* in rodents (most probable hosts in which hybridization occurs) [14], no other wild animals have been found with hybrid schistosomes. The situation in Corsica also needs to be precisely investigated.

### Implications for diagnostics

The hybrid status of the parasite may impair the parasitological, serological and molecular diagnostic. In endemic countries, parasitological diagnostic is the rule whereas serological tests are commonly used for imported schistosomiasis in non-endemic, developed countries. In humans, schistosome eggs that are partly retained in the tissues are the cause for the disease and induced pathology, but are also classical tools for diagnosis and species identification. At first sight, egg morphology and localization in the urine of infected patients in Corsica strongly suggested an *S. haematobium* infection [8]. Indeed *S. haematobium* eggs that are usually voided by the urine have a typical round to oval shape (elliptical or elongated) with a terminal spine. According to previous studies, *S. haematobium* eggs measure between 100-156 μm long and 40-50 μm wide with usual length between 115-135 μm long [15-17], A previous analysis of this Corsican hybrid schistosome eggs revealed smaller eggs (n=15) with a mean length of 106.5 μm and width of 42.8 μm, and with a spine length of 10.4 μm [13]. According to our results the eggs generally show an ovoid shape measuring 126.4 x 60.8 μm (Table 1) more similar to *S. haematobium* eggs. This is also consistent with the introgression levels that show a predominance of S. *haematobium-*type sequences (Table 3). Sometimes eggs were intermediate with spindle or diamond shapes, which are characteristic of *S. bovis eggs* (usually bigger and measuring between 170-223.9 μm long and 55-66.0 μm wide) and have an elongated spindle or diamond shapes [18,19] (Fig 1). In addition, whereas *S. haematobium eggs* are found in the urine of infected patients, *S. bovis* **eggs** are released in the feces. Thus, we can expect that for the hybrid parasite eggs may also be released in part in the feces. This could explain, together with the low parasite intensity, why only 30% of patients infected in Corsica excreted eggs in the urine [20]. The route of excretion associated with egg shape is the gold standard for diagnostic and species determination, nevertheless our results confirm earlier publications showing that it is impossible to detect hybridization in schistosome species using egg morphology alone [21].

Concerning the serological diagnostic, the majority of commercial tests, ELISA or IHA (indirect hemagglutination) use *S. mansoni* antigens. A discrepancy between those antigens and the infecting species may induce false negative results [22], The efficiency of these commercial diagnostic kits need thus to be reevaluated in a context of hybrid infection. Finally, molecular diagnostic for urogenital schistosomiasis using PCR has already been used in urine or serum, targeting a highly repeated sequence (DraI), which is restricted to the *S. haematobium* group of schistosomes (including both S. *haematobium* and *S. bovis*) [23,24], We expect that this test would be efficient to detect but not to identify the hybrid status of the parasite.

### Implications for treatment

Praziquantel (PZQ) is currently the only available drug used to treat schistosomiasis and the application of mass chemotherapy programs is the prevailing strategy for schistosomiasis control [25]. The treatment is efficient on bovine schistosomiasis but the dose needed (60 mg/kg for >95% deworming in goat [26,27]) cannot be considered in endemic areas were schistosomiasis is first of all a human health concern. Moreover, it has been shown that a dose of 40 mg/kg of PZQ is only 63.5% efficient for mixed infections, compared to 76.7% and 77.1%, for *S. mansoni* and *S. haematobium,* alone,respectively [28]. A lower sensitivity to PZQ of *S. bovis* x *S. haematobium* hybrid schistosomes compared to pure *S. haematobium* parasites has been proposed to be at the origin of the spread of the hybrid form in Senegal [29,30]. Hybridization may also affect Oxamniquine efficiency [31]. Since the genetic basis of Oxamniquine residence is known, our data can be used to propose a hypothesis on the degree of susceptibility. To date, neither experimental nor field trials have tested the sensitivity of hybrid parasites to praziquantel.

This work provides new insight into the schistosome hybrids that emerged in Europe, revealing admixture between *S. haematobium* and *S. bovis* parasites. As the strain we maintain in the lab has been recovered from a single infected patient in 2013, it is now necessary to extend our conclusions to subsequent contamination events in Corsica, and to investigate the dynamics of hybridization in the original endemic areas in Africa and especially in Senegal from where the parasite that have been introduced in Europe may have originated [8]. It is now essential to precisely characterize the impact of hybridization and introgression on the life history traits of the parasites, including sensitivity to current treatments, and assess the molecular mechanisms underlying these phenotypic changes. These hybrids may have the capacities to take over the geographical distribution of the parental species, but also rise out of endemic areas, and specifically in the close future with ongoing climate change, up north and towards Europe, making them an emerging global threat.

## Materials and Methods

### Ethics approval

Housing, feeding, animal care and experiments were carried out according to the national ethical standards established in the writ of 1 February 2013 (NOR: AGRG1238753A). The Direction Départementale de la Cohésion Sociale et de la Protection des Populations (DDSCPP) provided the permit N°C66-136-01 to our laboratory for experiments on animals. The investigator possesses the certificate for animal experimentation (Decree n° 87-848 du 19 octobre 1987; authorization 007083).

### Parasite / snail strains and experimental infections

The Corsican hybrid parasite was recovered from eggs found in the urine of an infected patient and maintained in the laboratory in sympatric snails *Bulinus truncatus* and hamsters *Mesocricetus auratus* [8]. Adult worms produced in hamsters were recovered after portal perfusion and male/female couples were manually separated. Detailed methods were described previously [32], *Schistosoma bovis* (Spain) and *Schistosoma haematobium* (Cameroon) were also maintained in the lab using *Planorbarius metidjensis* and *Bulinus truncatus* as intermediate hosts, respectively, and *Mesocricetus auratus* hamsters as definitive hosts [33]. F1 hybrids were produced after experimental cross between male *S. haematobium* and female *S. bovis.* For that, we infected molluscs with a single miracidium of the parental species to obtain male or female clonal cercariae. The latter were sexed using molecular markers efficient on both species that we developed previously [33]. We simultaneously exposed hamsters to 300 cercariae of male *S. haematobium* and 300 cercariae of female *S. bovis.* Three months after infection, hamsters were euthanized and F1 hybrid eggs were used to infect molluscs. The infected molluscs were then used to infest hamsters (pools of 600 cercariae) and adult male worms were collected 3 months after exposition.

### Hybrid parasite compatibility with snail hosts

*Planorbarius metidjensis* (n=40) and *Bulinus truncatus* (n=40) snails that are the natural hosts of *S. haematobium* and *S. bovis* respectively were individually exposed overnight to 5 miracidiae of the Corsican parasite strain maintained in the lab in well plates. Molluscs were individually checked from 35 days after infection for parasite emission of the cercariae after light stimulation.

### Morphological analysis of the Corsican schistosome hybrid eggs

Encysted eggs were collected from hamster livers and washed in 8.5% w/v Tris-NaCl. After whole-mounted on glass slides, eggs were viewed under light microscopy with objective lens magnification set to x10, and photographed on a Wild Heerbrugg M400 ZOOM Makroskop (Leica, Germany) coupled to Nikon digital sight DS - Fi1 digital camera. All measurements were produced with ImageJ version 1.51 [34] and drawings were done by image overlay in Adobe Photoshop CS2 version 9.0.1.

### DNA extraction and sequencing for *Schistosoma bovis* assembly

Two hundred clonal adult male worms were used to prepare high molecular weight genomic DNA using CHEF Genomic DNA Plug Kits (BioRad). The DNA was quantified on a FEMTO Pulse®, qubit and nanodrop. A total of 8.1 μg genomic DNA was used to generate a size selected PacBio library. First the DNA was sheared to an average fragment size of 45 kb by gently passing the DNA sample through a *2”* long, 26 gauge needle, 4 times and then concentrated using Ampure PB (Pacific Biosciences 100-265-900) before the library was prepared following the standard PacBio size selected library preparation protocol using BluePippin™ Size-Selection System. The library was size selected at 15kb, and run on 6 SMRT cells on the Sequel platform, generating 47.9 Gb of data.

### RNA extraction and sequencing for *Schistosoma bovis* annotation

Pools of 10-12 adult male or female worms were frozen with liquid nitrogen and grounded using Retsch MM400 cryobrush (2 pulses at 300Hz for 15s). Total RNA was extracted using TRIzol Thermo (Fisher Scientific) followed by DNase treatment with Turbo DNA-free kit. RNA was then purified using the RNeasy mini kit (Qiagen). The TruSeq stranded mRNA library construction kit (Illumina) was used on 300 ng of total RNA per condition. Library preparation and sequencing was performed at the McGill University in the Génome Québec Innovation Centre, Montréal, Canada on a Illumina HiSeq 4000 (100 bp paired-end reads).

### *Schistosoma bovis* genome assembly and annotation

A total of 48 Gb PacBio reads were assembled using HGAP4 to generate a 446 Mb assembly in 486 contigs, with an N50 of 3.1 Mb in 38 contigs.

Gene prediction was carried out with AUGUSTUS 3.3.1. No new training was performed for our data but the parameter set “schistosoma2” of the AUGUSTUS distribution was used. RepeatMasker 4.0.7 and RepeatScout 1.0.5 were used for repeat masking. RNAseq data from male and female adult worms was employed as external hints. This RNA-seq data was aligned with STAR version 020201.

### DNA extraction and sequencing for the Corsican hybrid strain and experimental *S*. *haematobium x S. bovis* F1

Genomic DNA of the Corsican schistosome hybrids was recovered from one pool of 10 adult males and one pool of 40 adult females separately, while DNA of the experimental F1 progeny was recovered from a pool of 10 adult males. DNA was extracted using Qiamp DNA microkit tissue kit (Qiagen) followed by RNase A treatment Genomic DNA of the Corsican hybrid worms was then sent to Genome Quebec for library construction using the Illumina TruSeq kit starting from 200 ng of genomic DNA (for each sex), and sequencing was performed on a Illumina HiSeq 2000 (100 bp paired-end reads). For the experimental F1 hybrid males, library construction was performed using the Nextera XT kit starting from 1 ng and sequenced on a Illumina NextSeq 550 (150 bp paired-end reads) on the Bio-Environment NGS platform at University of Perpignan.

### Estimation of the genomic introgression levels for the hybrid strain

The sequencing reads had PHRED quality scores over 30 with no adapter contamination. All reads were retained for further analysis and aligned to a chimeric concatenate of *S*. *haematobium* and *S. bovis* genomes using Bowtie 2 [35]. We used the SchistoDB *S*. *haematobium* genome [7] and the *S. bovis* SBOS_v1.1 assembly genome produced for this study. To avoid mapping bias due to differences in assembly size between the two genomes, only scaffolds > 1Mb were retained for further analysis. The genome size after concatenation of *S. haematobium* and *S. bovis* was 681.2 Mb. Mapping was thus performed by allowing each read, depending on its origin, to map against the more similar location in one or the other species genome. We then counted the proportion of best location of aligned reads (*S. bovis* or *S. haematobium* genomes) in the SAM files. The same procedure was applied on the mitochondrial genomes using a concatenate of the scaffold 000439F that contained the mitochondrial genome of *S. bovis,* and the *S. haematobium* mitochondrial genome from GenBank accession NC_008074.

### Similarity analysis between *S. haematobium* and *S. bovis* genomes

The two species genomes were aligned with CACTUS [36]. We then processed the output, identifying all alignments blocks composed of exactly one genome part of *S. bovis* and exactly one genome part of *S. haematobium.*

### Availability of data

The Illumina datasets for the Corsican hybrid strain are available in the SRA repository under submission number SUB4330327 (to be released upon publication). The *S. bovis* PacBio data are available under accession number ERS2549235. The *Schistosoma bovis* assembly SBOS_vl.l is available at ftp://ftp.sanger.ac.uk/pub/project/pathogens/Schistosoma/bovis/SBOS_vl.l_pilon.fast a. Annotations are available as trackhub at http://ihpe.univ-perp.fr/IHPE_tracks/S.bovis/hub.txt

## Acknowledgments

We would like to acknowledge the NGS facility at the Bio-Environment platform (University of Perpignan) and DNA Pipelines at Wellcome Sanger Institute, in particular Michelle Smith and Craig Corton.

## Authors’ contributions

ET and JB conceived and designed the study. JB, AO and SMC procured the parasite and snail strains. JKS, AR, and JB performed the experiments. RA conducted the morphological analysis. NH and JFA prepared the DNA for sequencing. NH coordinated the long read DNA sequencing experiments. JKS, AT, IB, MB and ET conducted the computational data processing and analysis with significant assistance and input from OR, CC and CG. JKS and ET wrote the paper. All authors read and approved the final manuscript.

## Supporting information captions

**Appendix S1: Morphological analysis of the European schistosome hybrid eggs and adult worms**

## References

1. Chitsulo L, Chitsulo L, Engels D, Engels D, Montresor A, Montresor A, et al. The global status of schistosomiasis and its control. Acta Trop. 2000;77: 41–51. doi: 10.1016/s0001-706x(00)00122-4

2. De Bont J, Vercruysse J. The epidemiology and control of cattle schistosomiasis. Parasitol Today. 1997;13: 255–262.

3. Wu X, Lu Y, Zhou S, Chen L, Xu B. Impact of climate change on human infectious diseases: Empirical evidence and human adaptation. Environ Int. 2016;86: 14–23. doi: 10.1016/j.envint.2015.09.007

4. Bar-Zvi D, Lupo O, Levy AA, Barkai N. Hybrid vigor: The best of both parents, osr a genomic clash? Current Opinion in Systems Biology. 2017;6: 22–27. doi: 10.1016/j.coisb.2017.08.004

5. King KC, Stelkens RB, Webster JP, Smith DF, Brockhurst MA. Hybridization in Parasites: Consequences for Adaptive Evolution, Pathogenesis, and Public Health in a Changing World. PLoS Pathog. 2015;11: el005098. doi:10.1371/journal.ppatl005098.g001

6. Léger E, Webster JP. Hybridizations within the Genus Schistosoma: implications for evolution, epidemiology and control. Parasitology. 2017.

7. Young ND, Jex AR, Li B, Liu S, Yang L, Xiong Z, et al. Whole-genome sequence of Schistosoma haematobium. Nat Genet 2012;44: 221–225. doi:10.1038/ng,1065

8. Boissier J, Grech-Angelini S, Webster BL, Allienne J-F, Huyse T, Mas-Coma S, et al. Outbreak of urogenital schistosomiasis in Corsica (France): an epidemiological case study. The Lancet Infectious Diseases. 2016;16: 971–979. doi: 10.1016/S1473-3099(16)00175-4

9. Valentim CLL, Cioli D, Chevalier FD, Cao X, Taylor AB, Holloway SP, et al. Genetic and Molecular Basis of Drug Resistance and Species-Specific Drug Action in Schistosome Parasites. Science. 2013;342:1385–1389. doi:10.1093/molbev/msrl21

10. Chevalier FD, Le Clec’h W, Eng N, Rugel AR, Assis RR de, Oliveira G, et al. Independent origins of loss-of-function mutations conferring oxamniquine resistance in a Brazilian schistosome population. International Journal for Parasitology. 2016;46: 417–424. doi:10.1016/j.ijpara.2016.03.006

11. Detwiler JT, Criscione CD. An infectious topic in reticulate evolution: introgression and hybridization in animal parasites. Genes (Basel). 2010;1: 102–123. doi:10.3390/genesl010102

12. Holtfreter MC, Mone H, Miiller-Stover I, Mouahid G, Richter J. Schistosoma haematobium infections acquired in Corsica, France, August 2013. Euro Surveill. 2014;19.

13. Moné H, Holtfreter MC, Allienne J-F, Mintsa-Nguéma R, Ibikounlé M, Boissier J, et al. Introgressive hybridizations of Schistosoma haematobium by Schistosoma bovis at the origin of the first case report of schistosomiasis in Corsica (France, Europe). Parasitol Res. 2015. doi:10.1007/s00436-015-4643-4

14. Catalano S, Séne M, Diouf ND, Fall CB, Borlase A, Leger E, et al. Rodents as Natural Hosts of Zoonotic Schistosoma Species and Hybrids: An Epidemiological and Evolutionary Perspective From West Africa. J Infect Dis. 2018;218: 429–433. doi:10.1093/infdis/jiy029

15. Moné H, Minguez S, Ibikounlé M, Allienne J-F, Massougbodji A, Mouahid G. Natural Interactions between S. haematobium and S. guineensis in the Republic of Benin. ScientificWorldJournal. 2012;2012: 793420. doi:10.1100/2012/793420

16. Loker ES. A comparative study of the life-histories of mammalian schistosomes. Parasitology. 1983;87 (Pt2): 343–369.

17. Pitchford RJ. Differences in the egg morphology and certain biological characteristics of some African and Middle Eastern schistosomes, genus Schistosoma, with terminal-spined eggs. Bull World Health Organ. 1965;32:105-120.

18. Rollinson D, Southgate VR, Vercruysse J, Moore PJ. Observations on natural and experimental interactions between Schistosoma bovis and S. curassoni from West Africa. Acta Trop. 1990;47: 101–114.

19. Alves W. The eggs of Schistosoma bovis, S. mattheei and S. haematobium. J Helminthol. 1949;23: 127–134.

20. Noël H, Ruello M, Maccary A, Pelat C, Sommen C, Boissier J, et al. Large outbreak of urogenital schistosomiasis acquired in Southern Corsica, France: monitoring early signs of endemicization? Clin Microbiol Infect 2018;24: 295–300. doi:10.1016/j.cmi.2017.06.026

21. Boon N, Fannes W, Rombouts S, Polman K, Volckaert FAM, Huyse T. Detecting hybridization in African schistosome species: does egg morphology complement molecular species identification? Parasitology. 2017;144: 954–964. doi:10.1017/S0031182017000087

22. Kinkel H-F, Dittrich S, Bäumer B, Weitzel T. Evaluation of eight serological tests for diagnosis of imported schistosomiasis. Clin Vaccine Immunol. 2012;19: 948–953. doi:10.1128/CVI.05680-ll

23. Hamburger J He-Na, Abbasi I, Ramzy RM, Jourdane J, Ruppel A. Polymerase chain reaction assay based on a highly repeated sequence of Schistosoma haematobium: a potential tool for monitoring schistosome-infested water. Am J Trop Med Hyg. 2001;65: 907–911.

24. Cnops L, Soentjens P, Clerinx J, Van Esbroeck M. A Schistosoma haematobium-specific real-time PCR for diagnosis of urogenital schistosomiasis in serum samples of international travelers and migrants. PLoS Negl Trop Dis. 2013;7: e2413. doi:10.1371/journal.pntd.0002413

25. Colley DG, Bustinduy AL, Secor WE, King CH. Human schistosomiasis. Lancet. 2014;383: 2253–2264. doi:10.1016/S0140-6736(13)61949-2

26. Monrad J, Sörén K, Johansen MV, Lindberg R, Ornbjerg N. Treatment efficacy and regulatory host responses in chronic experimental Schistosoma bovis infections in goats. Parasitology. 2006;133:151–158. doi:10.1017/S0031182006000102

27. Johansen MV, Monrad J, Christensen NO. Effects of praziquantel on experimental Schistosoma bovis infection in goats. Vet Parasitol. 1996;62: 83–91.

28. Zwang J, Olliaro PL. Clinical efficacy and tolerability of praziquantel for intestinal and urinary schistosomiasis-a meta-analysis of comparative and non-comparative clinical trials. PLoS Negl Trop Dis. 2014;8: e3286. doi:10.1371/journal.pntd.0003286

29. Huyse T, Webster BL, Geldof S, Stothard JR, Diaw OT, Polman K, et al. Bidirectional introgressive hybridization between a cattle and human schistosome species. PLoS Pathog. 2009;5: el000571–el000571. doi:10.1371/journal.ppatl000571

30. Webster BL, Diaw OT, Seye MM, Webster JP, Rollinson D. Introgressive hybridization of Schistosoma haematobium group species in Senegal: species barrier break down between ruminant and human schistosomes. PLoS Negl Trop Dis. 2013;7: e2110. doi:10.1371/journal.pntd.0002110

31. Pitchford RJ, Lewis M. Oxamniquine in the treatment of various schistosome infections in South Africa. S Afr Med J. 1978;53: 677–680.

32. Boissier J, Chlichlia K, Digon Y, Ruppel A, Moné H. Preliminary study on sex-related inflammatory reactions in mice infected with Schistosoma mansoni. Parasitol Res. 2003;91:144–150. doi:10.1007/s00436-003-0943-l

33. Kincaid-Smith J, Boissier J, Allienne J-F, Oleaga A, Djuikwo-Teukeng F, Toulza E. A Genome Wide Comparison to Identify Markers to Differentiate the Sex of Larval Stages of Schistosoma haematobium, Schistosoma bovis and their Respective Hybrids. PLoS Negl Trop Dis. 2016;10: e0005138. doi:10.1371/journal.pntd.0005138

34. Prodanov D, Verstreken K. Automated Segmentation and Morphometry of Cell and Tissue Structures. Selected Algorithms in ImageJ. 2012.

35. Langmead B, Langmead B, Trapnell C, Trapnell C, Pop M, Pop M, et al. Ultrafast and memory-efficient alignment of short DNA sequences to the human genome. Genome Biol. 2009;10. doi:10.1186/gb-2009-10-3-r25

36. Paten B, Earl D, Nguyen N, Diekhans M, Zerbino D, Haussler D. Cactus: Algorithms for genome multiple sequence alignment Genome Research. 2011;21: 1512–1528. doi:10.1101/gr.123356. III

