## Appendix S1 for "Morphological and genomic characterisation of the hybrid schistosome infecting humans in Europe reveals a complex admixture between *Schistosoma haematobium and Schistosoma bovis parasites*"

### Appendix S1: Morphological analysis of the European schistosome hybrid eggs and adult worms

#### Material and methods

Hamsters infected with the Corsican hybrids schistosomes were euthanized and the encysted eggs accumulated in hamster’s tissues (liver) were collected in 8,5% NACL solution to analyse their morphology. Eggs were put on glass slides and viewed under light microscopy with objective lens magnification of x10. We measured the length (including the spine), the width (at it’s larger point), the size of the terminal spine, and the total area of the eggs using the open-source imaging program ImageJ.

At the same time 20 adult couples (20 male and 20 female) of worms were fixed and intended to morphological description. Adults’ worms were collected from hamster’s hepatic perfusion, and washed in 8,5% Tris-NaCl solution before manually separating males and females and storing them in 70% ethanol. For light microscopy, the schistosomes adults were whole-mounted on glass slides. Adult worms and eggs were photographed on a Wild Heerbrugg M400 ZOOM Makroskop (Leica, Germany) or Dialux20 (Leitz, Germany) coupled to Nikon digital sight DS – Fi1 digital camera and all measurements were given in micrometers (mm) with ImageJ version 1.51. The following characters were measured in worms of both sexes: worm length and width, area of the oral and ventral suckers, sucker’s ratio, sucker ratio per worm length, distance from genital opening to anterior region. In female worms the area of the ovary area and extension of the vitellines were measured. In males we also recorded the number of testes. Drawings were done by image overlay in Adobe Photoshop CS2 version 9.0.1.

#### Results

A total of 44 eggs were examined for morphological characterization. The length and width were assessed on all eggs, but the spine lengths have been measured on a subset of 36 that had a spine distinctive enough to allow for proper estimation of their sizes. The results are presented in Table 1. Hybrid eggs had a mean length of 126.4 µm (±22.9 µm SD), and a mean width of 60.8 µm (±13.0 µm SD). The mean spine length was of 8.2 µm (±2.1 µm SD). The minimum-maximum and number of eggs analyzed for each variable are presented in Table 1.

Table 1: Morphological measurements results of the Corsican hybrid schistosome eggs (SD = Standard deviation).

|  | **Length**  **(including spine)** | **Width** | **Spine length** |
| --- | --- | --- | --- |
| **No. of eggs** | **44** | **44** | **36** |
| **Mean (**± SD) in **µm** | 126.4 (±22.9) | 60.8 (±13.0) | 8.2 (±2.1) |
| **Minimum** in **µm** | 73.9 | 40.9 | 3.95 |
| **Maximum** in **µm** | 170.9 | 92.5 | 13.6 |

Most of the eggs displayed a representative elliptical morphotypes and were characterized by a terminal spine, reminiscent of *S. haematobium* infection in humans. Nevertheless, not all eggs had a typical *S. haematobium* morphotype and in some cases they seemed intermediate with *S. bovis*-type eggs (Fig 1).

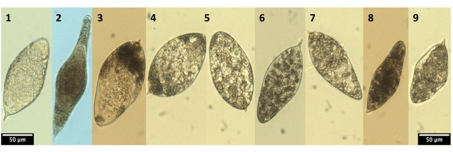

**Figure 1: Egg morphologies of the pure parental species and the Corsican *S. haematobium X S. bovis* hybrid**. Eggs 1 and 2 show typical morphologies of *S. haematobium* (elliptical with a terminal spine) and *S. bovis* (spindle shape with a terminal spine), respectively. Eggs 3-8 show the egg morphology of the Corsican hybrid schistosome. While most eggs were typical to *S. haematobium* (3-7), a high variability of morphotypes was observed (8-9).

Like other species of the genus *Schistosoma*, male and female adults of the schistosome hybrids exhibited a characteristic marked sexual dimorphism and females living in the gynaecophoric canal of male worms (Fig 2). The detailed morphological characteristic measurements are presented in Table 2.

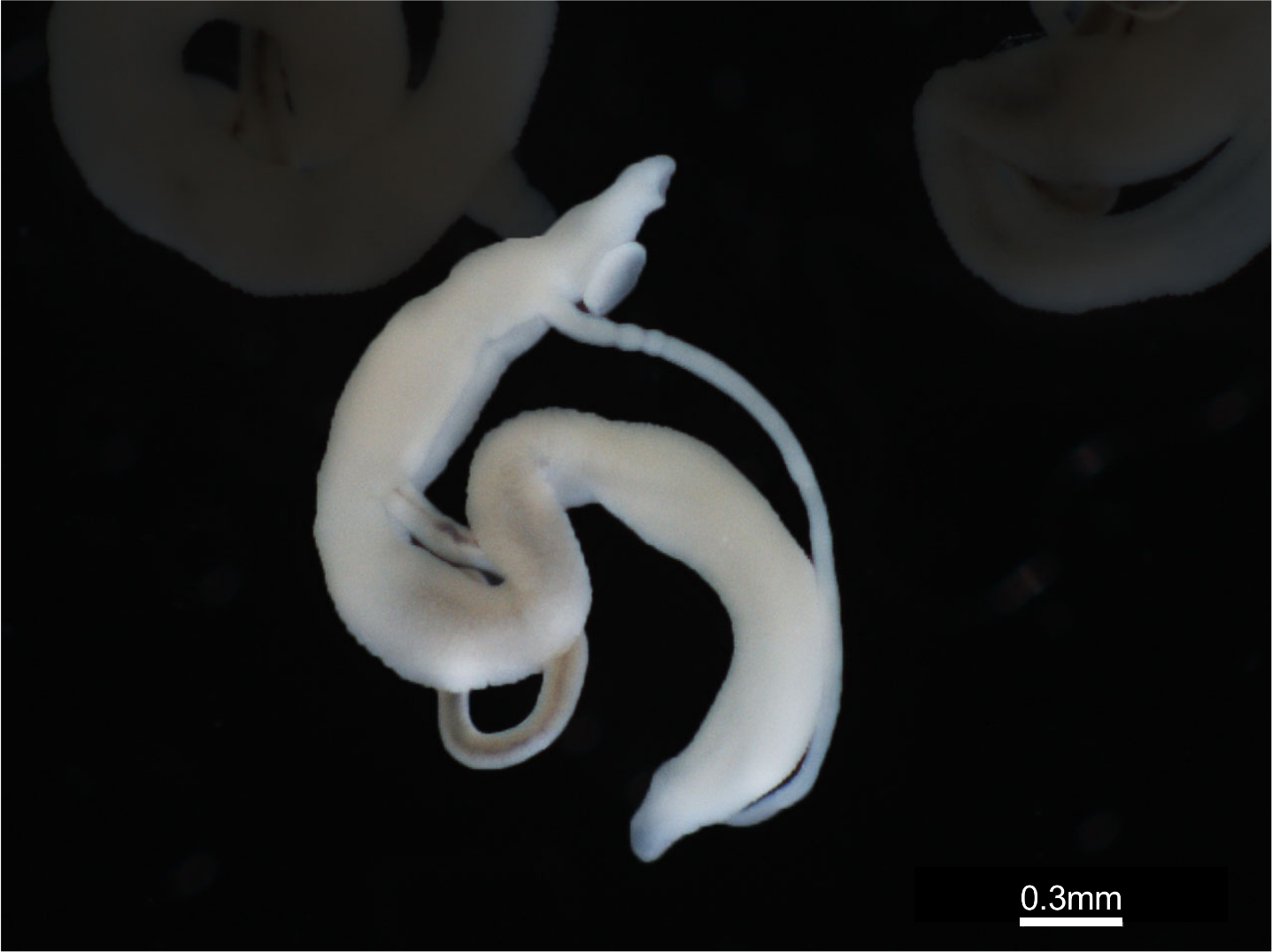

**Figure 2: Morphology of natural hybrid from Corsican strain.** Sexual dimorphism of the hybrid schistosomes with adult females living in the gynaecophoric canal of adult male parasites.

**Table 2: Morphological measurements of adult Corsican hybrid male and female schistosomes**

|  | | **Hybrid schistosomes from Corsica** | | | |
| --- | --- | --- | --- | --- | --- |
|  |  | **Male** | | **Female** | |
|  |  | **Mean (**±**SD)** | **Range** | **Mean (**±**SD)** | **Range** |
| **Total length (mm)** | 5.3 (±1.0) | | 3.3 – 6.6 | 9.6 (±1.2) | 8.6 – 14.1 |
| **Largest body width (mm)** | 0.3 (±0.07) | | 0.2 – 0.5 | 0.2 (±0.04) | 0.1 – 0.3 |
| **Oral sucker length (mm)** | 0.2 (±0.01) | | 0.2 – 0.2 | 0.05 (±0.004) | 0.04 – 0.06 |
| **Oral sucker width (mm)** | 0.2 (±0.006) | | 0.1 – 0.2 | 0.05 (±0.002) | 0.05 – 0.06 |
| **Ventral sucker length (mm)** | 0.3 (±0.008) | | 0.2 – 0.3 | 0.06 (±0.001) | 0.06 – 0.07 |
| **Ventral sucker width (mm)** | 0.3 (±0.01) | | 0.2 – 0.3 | 0.06 (±0.001) | 0.06 – 0.07 |
| **Testes number** | 4.2 (±0.5) | | 4 –5 | - | - |
| **Ovary length (mm)** | - | | - | 0.3 (±0.01) | 0.3 – 0.3 |
| **Ovary width (mm)** | - | | - | 0.09 (±0.01) | 0.08 – 0.1 |
| **Extension of the vitellaria (mm)** | - | | - | 5.0 (±0.7) | 4 – 5.9 |

The adult male parasites had an elongated body measuring 5.3 (±1.0) mm long and 0.3 (±0.07) mm wide at largest part of the body. The male body was dorso-ventrally flattened with folds in the ventral axis forming the gynaecophoral canal where the paired females are commonly found (Fig 3a). The anterior region was narrower than the rest of the body and presented a sub-terminal mouth formed into the oral sucker (length: 0.2±0.01 mm; width: 0.2±0.006 mm) and a robust ventral sucker or acetabulum (length: 0.3±0.008 mm; width: 0.3±0.01 mm) near the top of the gynaecophoral canal (Fig 3b). The oesophagus began at the oral sucker and was extended posteriorly to the acetabulum where it bifurcates into the gut caeca until it reunites at the posterior end of the body.

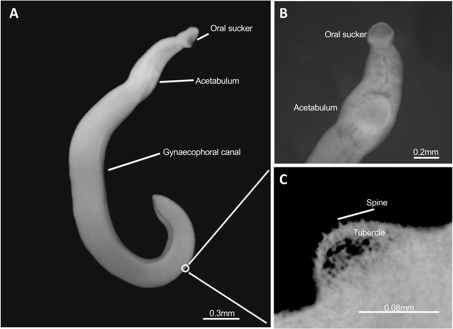

Figure 3: Morphological observations of male hybrid schistosomes from Corsica: (A) Whole parasite body locating oral sucker, acetabulum (ventral sucker) and gynaecophoral canal; (B) Frontal view of anterior region showing oral sucker, acetabulum (ventral sucker) in detail; (C) Dorso-lateral view of spines and tubercle over parasite surface in detail.

The testes, situated dorsally posterior to the ventral sucker, were round to ovoid and we found 4 to 5 in number in adult males (4.2±0.5). Posterior to the acetabulum, in the dorsal region, tubercles with a round extremity began to appear at the level of the gynaecophorial canal and occurred to all the posterior region of the body (Fig 3c). The presence of tegument projections on the tubercles where identified with several apical spines which decreased in distribution and size towards the back and sides of the male’s bodies (Fig 3c). The female’s bodies were elongated and filiform and measured 9.6 (±1.2) mm long and 0.2 (±0.04) mm wide, with the posterior half of body expanded (Fig 4a). The anterior regions were smaller compared to males, and females had a small oral sucker (length: 0.05±0.004 mm; width: 0.05±0.002 mm) and acetabulum (length: 0.06±0.001 mm; width: 0.06±0.001 mm) (Fig 4b).

The oesophagus started near the oral sucker and bifurcated immediately after the acetabulum. The genital pore was situated at the posterior of the ventral sucker (dorsally) and some eggs could be observed in the uterus (Fig.4c). A single ovary measuring 0.3 (±0.01) long and 0.09 (±0.01) wide was situated in the posterior third of the female’s body (Fig 4a). The vitelline glands, also called vitellaria was extensive, occupying roughly 50% of the posterior part of the worm and extending further posteriorly than the intestine. The females’ tegument was smooth and uniform without significant projections. The posterior extremity was tapered and rounded.

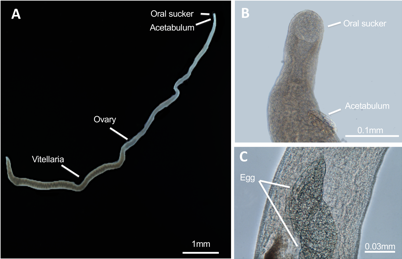

**Figure 4: Morphological observations of female hybrid schistosomes from Corsica:** (A) Lateral view of whole parasite body showing oral sucker, acetabulum (ventral sucker), ovary and vitellaria; (B) Frontal view of anterior region showing oral sucker, acetabulum (ventral sucker) in detail; (C) Dorso-lateral view of two eggs inside female uterus in detail.
